# STAT5 regulation of sex-dependent hepatic CpG methylation at distal regulatory elements mapping to sex-biased genes

**DOI:** 10.1101/2020.04.22.054601

**Authors:** Pengying Hao, David J. Waxman

**Affiliations:** Department of Biology and Bioinformatics Program, Boston University, Boston, MA 02215

**Keywords:** RRBS, enhancers, CpG methylation, sex-bias, DNase hypersensitive site

## Abstract

Growth hormone-activated STAT5b is an essential regulator of sex-differential gene expression in mouse liver, however, its impact on hepatic gene expression and epigenetic responses is poorly understood. Here, we found a substantial, albeit incomplete loss of liver sex bias in hepatocyte-specific STAT5a/STAT5b (collectively, STAT5)-deficient mouse liver. In male liver, many male-biased genes were down regulated in direct association with the loss of STAT5 binding; many female-biased genes, which show low STAT5 binding, were de-repressed, indicating an indirect mechanism for repression by STAT5. Extensive changes in CpG-methylation were seen in STAT5-deficient liver, where sex differences in DNA methylation were abolished at 88% of ~1,500 differentially-methylated regions, largely due to an increase in methylation at the hypomethylated sites. STAT5-dependent CpG-hypomethylation was rarely found at proximal promoters of STAT5-dependent genes. Rather, STAT5 primarily regulated the methylation of distal enhancers, where STAT5 deficiency induced widespread hypermethylation at genomic regions enriched for accessible chromatin, enhancer histone marks (H3K4me1, H3K27ac), STAT5 binding, and DNA motifs for STAT5 and other transcription factors implicated in liver sex differences. In conclusion, the sex-dependent binding of STAT5 to liver chromatin is closely linked to sex-dependent demethylation of distal regulatory elements mapping to STAT5-dependent genes important for liver sex bias.

## Introduction

The mammalian STAT gene family encodes seven latent cytoplasmic transcription factors activated by cell surface receptor-induced tyrosine phosphorylation, which stimulates STAT protein dimerization and nuclear translocation, followed by DNA binding and transcriptional activation of STAT target genes (1). One STAT gene, STAT5a, is highly expressed in mammary tissues, where it is activated by prolactin (2, 3), while the closely related (96% similar) STAT5b is particularly abundant in hepatocytes, where it is activated by growth hormone (GH) and regulates lipid metabolism (4–6) as well as body growth, in part through production of IGF-1 (7). STAT5b is also a key mediator of the sex differential transcriptional networks that GH regulates in the liver (8, 9), as seen in a STAT5b whole body knockout mouse model (10). GH activates STAT5b signaling by binding to its liver cell surface receptor, which in turn activates the receptor-associated tyrosine kinase JAK2, stimulating phosphorylation of GH receptor on multiples intracellular tyrosine residues, thereby creating docking sites for downstream signaling proteins, including STAT5b. JAK2-catalyzed phosphorylation of STAT5b on Tyr-699 enables STAT5b to dimerize and then translocate to the nucleus (11), where it binds to STAT5 motifs enriched at open chromatin regions and stimulates gene transcription (12).

Liver STAT5 activity is highly responsive to GH stimulation (13). In many mammalian species, including humans (14), rats and mice (8, 15), the temporal pattern of pituitary GH secretion differs between males and females. In male mice and rats, pituitary GH secretion is episodic, with strong pulses every 3-4 h followed by a GH-free interval, whereas in females, pituitary GH secretion is more frequent, resulting in a persistent (near continuous) plasma GH profile. Liver STAT5 activity mirrors the sex differences in circulating GH patterns, and consequently, it oscillates between active and inactive states in male liver, but is more persistently active in female liver (12, 16, 17). These sex-dependent temporal differences in liver STAT5 activity are the dominant determinant of the sex-biased expression of hundreds of genes in mouse and rat liver, including many cytochromes P450 and other enzymes of steroid and drug metabolism, as was demonstrated in mice with a global deficiency in STAT5b (10), and are an important factor in sex-differences in drug and steroid metabolism and disease susceptibility (8, 18). STAT5a deficiency can also impact sex-biased gene expression in the liver, albeit to a much lesser extent than STAT5b, consistent with the much lower levels of STAT5a present in liver (19). Perturbation of liver STAT5 activity by altering plasma GH patterns by hypophysectomy (16, 20) or by continuous GH infusion (21, 22) largely abolishes sex differences in gene expression, including the expression of sex-dependent regulatory micro-RNAs (23) and lncRNAs (24, 25). Importantly, while exogenous GH replacement can restore sex-specific expression of *Cyp* and other genes in livers of hypophysectomized mice, this restoration is not achieved in livers of hypophysectomized STAT5b knockout mice (22, 26). While these studies demonstrate the essential nature of STAT5b for sex differentiated liver gene expression, they do not distinguish effects due to the loss of STAT5b in liver *per se* versus effects due to loss of STAT5 expression in other tissues, including the hypothalamus, which can impact plasma GH profiles and liver responses to GH (27). These issues are addressed here, where we use RNA-seq to evaluate the impact of hepatocyte-specific deletion of the *Stat5a/Stat5b* locus (28) on the liver transcriptome in both male and female mice, and its relationship to sex-specific binding sites for STAT5 (12, 29) associated with regulation of sex-biased gene expression in the liver.

DNA methyltransferase-catalyzed cytosine methylation at the C-5 position is a heritable epigenetic mark in higher vertebrates and is key for many biological processes, including X chromosome inactivation and imprinting (30, 31). A large majority of DNA methylation occurs in the context of CpG dinucleotides in mammalian somatic cells, with most CpG sites being highly methylated, except for CpGs within gene promoters. These patterns of DNA methylation are stably propagated in terminally differentiated cells throughout life; however, changes in CpG methylation can occur in response to hormonal and other environmental factors (32, 33) and diseased states (30). DNA methylation generally leads to gene silencing and chromatin condensation, although DNA methylation can also stimulate gene expression (34). Sex-differences in DNA methylation have been reported for mouse liver (35–37) and may, in part, be determined by CpG demethylation associated with testosterone exposure at puberty (38–40). It is unclear, however, what role STAT5 might play in regulating these sex differences in CpG methylation, or how they may relate to the STAT5-dependent regulation of sex-specific gene expression. Here, we use reduced representation bisulfite sequencing (41) to characterize the effects of STAT5 on CpG methylation, including sex differences in liver CpG methylation. Our findings establish that liver STAT5 deficiency abolishes the sex bias in DNA methylation at more than 1,300 genomic sites, and that STAT5-dependent sex-biased genes are frequently regulated by STAT5-regulated demethylation associated with the activation of distal enhancers.

## Results

### Impact of hepatocyte-specific STAT5 loss on sex-biased liver gene expression

Global inactivation of the mouse *Stat5b* gene leads to widespread loss of sex-biased gene expression in the liver (10). As the observed changes in liver gene expression could, in part, be due to extrahepatic actions of STAT5b on pituitary GH secretion (26), which is a major regular of sex-specific gene expression in the liver (21), we used RNA-seq to determine the impact of hepatocyte-specific deletion of the *Stat5a/Stat5b* locus (STAT5-KO) (28, 42) on sex-biased gene expression. Differential expression analysis was carried out using liver RNA purified from wild-type male (WTM) and female (WTF) floxed control mice, and from STAT5-KO male (KOM) and female (KOF) mice. A total of 274 liver-expressed genes showed differential expression between male and female controls (WTM/WTF comparison, FDR < 0.05): 132 male-biased and 142 female-biased genes. Genes that were responsive to STAT5-KO in male liver (365 genes) and in female liver (278 genes) were identified in two additional comparisons, KOM/WTM and KOF/WTF (Table S1). The gene overlap across the three comparisons is presented in **Fig. 1A**. 155 of all 274 sex-biased genes (57%) responded to STAT5 deficiency in either male or female liver vs. only 2.2% of stringently sex-independent genes (**Table 1**). Male-biased genes whose expression was altered were primarily down regulated in STAT5-KO male liver or were up regulated in STAT5-KO female liver. Conversely, female-biased genes whose expression was altered were primarily up regulated in STAT5-KO male liver or were down regulated in STAT5-KO female liver (**Fig. 1B** and **Table 1**). This relationship between sex specificity and response to STAT5 deficiency is visualized in quantitative scatter plots showing highly significant differences in the response of male-biased and female-biased genes to STAT5-KO loss in male liver, and to a lesser degree in female liver (**Fig. 1C**). The extent of sex bias and magnitude of response to STAT5 deficiency were significantly correlated in male liver (r^2^=0.4, p = 1.2E-13), but not in female liver. Thus, STAT5 acts at the level of the hepatocyte to regulate sex-biased gene expression *via* both activating and repressive mechanisms, strongly in male liver and weakly in female liver.

**Fig. 1 –.**
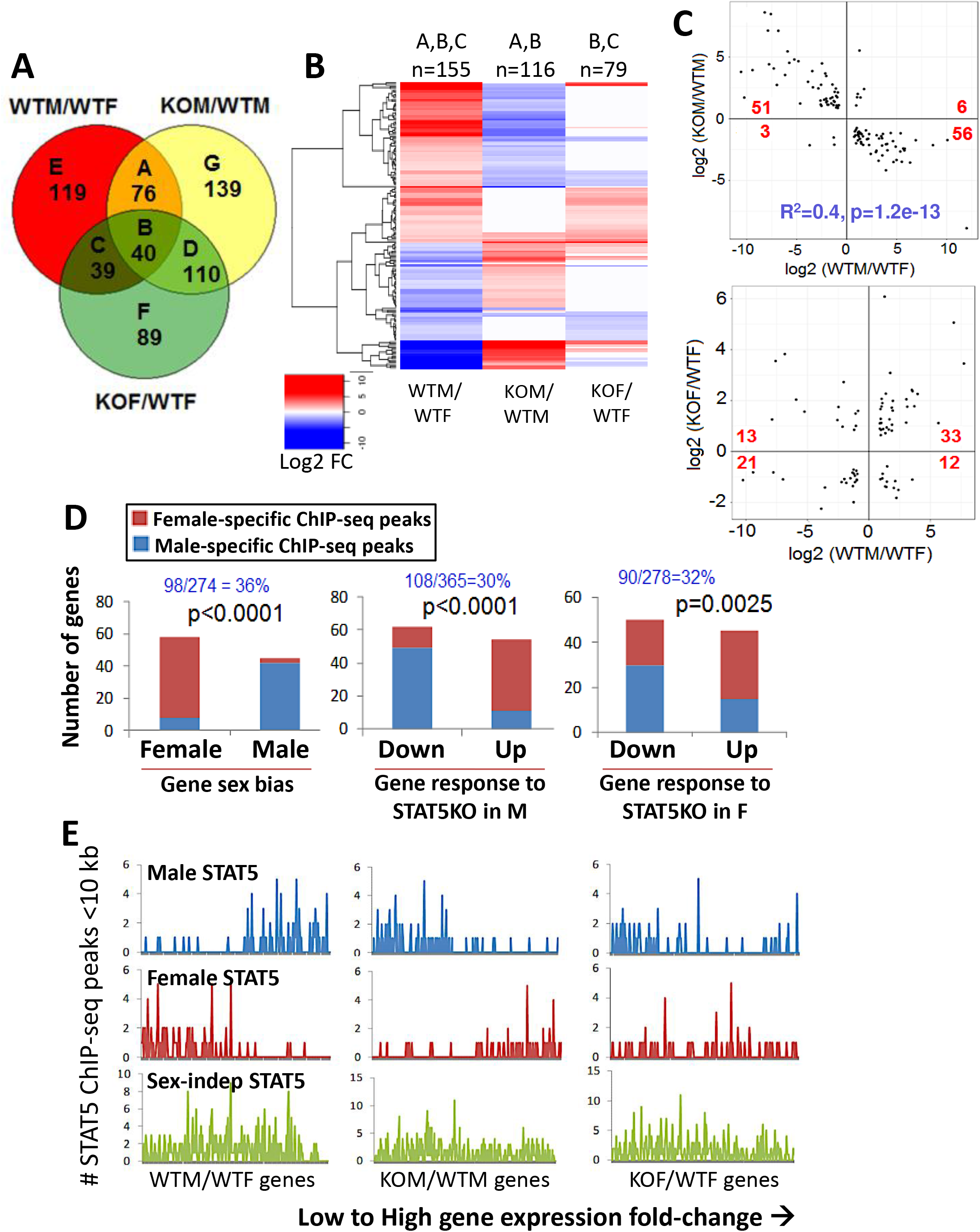
Gene expression changes in STAT5a/STAT5b-deficient mouse liver. (**A**) Venn diagram showing 612 significantly changing genes (FDR<0.05) identified in wild-type (WT) and STAT5 knockout (KO) male (M) and female (F) mouse liver for three comparisons: WTM/WTF (wild-type male versus wild-type female; sex-biased genes); KOM/WTM (knockout male versus wild-type male; STAT5 responsive genes in male liver) and KOF/WTF (knockout female versus wild-type female; STAT5 response genes in female liver). The number of differentially expressed genes in each comparison (labeled A through G) is indicated. (**B**) Heat map representation of log2 fold-change of 155 genes that showed sex specific expression in wild-type liver (WTM/WTF) and their corresponding changes in STAT5-KO male (KOM/WTM) and STAT5-KO female (KOF/WTF) livers. These genes correspond to regions A, B and C in the Venn diagram, as indicated at the top. n, number of genes. (**C**) Scatter plots of log2 expression fold-change values of sex-biased genes (WTM/WTF) that responded significantly to STAT5 loss in male liver (116 genes; regions A and B of Venn diagram; top), or that responded significantly to STAT5 loss in female liver (79 genes; regions B, C of Venn diagram; bottom). The number of genes in each quadrant of the plot was analyzed by Fisher’s exact test to assess whether the proportions of male and female biased genes that were up regulated and down regulated by STAT5 loss are significantly different: p<2.2E-16 (*top*) and p=0.003 (*bottom*). Linear regression was performed on the absolute values of fold change values to determine the level of correlation between magnitude of sex bias and strength of response to STAT5 deletion: r^2^=0.40, p=1.2E-13 (top), and r^2^=0.07, p=0.01 (bottom). (**D**) Relationship between STAT5 transcription factor binding and liver gene expression. Genes with one or more sex-biased STAT5 ChIP-seq peaks within 10 kb were counted (y-axis) and is indicated in blue text as a fraction of all genes in each gene set (*top*). (*Left*) In wild-type liver, female-biased genes were significantly enriched for having nearby female-biased STAT5 ChIP-seq peaks, and male-biased genes for having nearby male-biased STAT5 ChIP-seq peaks. (*Middle*) In male liver, genes down regulated by STAT5-KO were significantly enriched for nearby male-biased STAT5 ChIP-seq peaks, and genes up regulated by STAT5-KO were enriched for nearby female-biased STAT5 ChIP-seq peaks. P-values for the sex-difference in these response patterns are shown (Fisher’s exact test). (*Right*) In female liver, STAT5 responsive genes followed the same pattern as those in male liver, albeit with less significant differential enrichment for sex-biased STAT5 ChIP-seq peaks (higher p-value). (**E**) Shown are the number of STAT5 ChIP-seq peaks (male-biased, *top* row of graphs, in blue; female-biased, *middle* row, in red; or sex-independent, *bottom* row, in green) within 10 kb of each individual gene that is differentially expressed at FDR<0.05 between: WTM/WTF (274 genes, *left*), KOM/WTM (365 genes, *middle*) and KOF/WTF (278 genes, *right*), ranked from low to high gene expression fold-change. This represents a per gene visual representation of the results calculated in (D). See Table S1 for detailed listing of expression values for these 612 genes.

**Table 1.** Liver-expressed genes showing significant expression changes in hepatocyte specific STAT5-KO mouse liver. 414 liver-expressed genes (FPKM >1 in either wild-type male or female liver) were analyzed: 132 male-biased genes, 142 female-biased genes, and 140 stringently sex-independent genes whose expression changes significantly (FDR < 0.05) with STAT5 deficiency in either male or female mouse liver (Table S1A, Table S1B). Data are shown for the 155 of 274 sex-biased genes (57%; 87 of 132 male-biased genes and 68 of 142 female-biased genes) and for the 140 of 6,323 stringently sex-independent genes (2.2%) that are liver-expressed and whose expression changes significantly with STAT5 deficiency in either male or female mouse liver (genes ‘responsive’ to STAT5-KO). Percentage values indicate the distribution of each gene group between up and down regulation in STAT5-deficient mouse liver. Response patterns to STAT5 deficiency were significantly different by Fisher’s Exact test: between male-biased and female-biased genes, and between female-biased and sex-independent genes, in male livers, and also in female livers; and between male-biased and sex-independent genes in male liver only. Response patterns were also significantly different for male-biased genes between male and female livers, and for female-biased genes between male and female livers.

| Sex-bias in wild-type liver | Fraction of KO-responsive genes | Change in expression in STAT5-KO | Response to STAT5-KO in male liver |  | Response to STAT5-KO in female liver |  |
| --- | --- | --- | --- | --- | --- | --- |
|  |  |  | Gene count | % | Gene count | % |
| Male-biased and KO-responsive | (87/132 = 66%) | Increase | 6 | 9.7 | 33 | 73.3 |
|  |  | Decrease | 56 | 90.3 | 12 | 26.7 |
| Female-biased and KO-responsive | (68/142 = 48%) | Increase | 51 | 94.4 | 13 | 38.2 |
|  |  | Decrease | 3 | 5.6 | 21 | 61.8 |
| Sex-independent and KO-responsive | (140/6323 = 2.2%) | Increase | 65 | 61.3 | 58 | 66.7 |
|  |  | Decrease | 41 | 38.7 | 29 | 33.3 |

Sex-biased expression was lost in STAT5-KO liver for 168 (61%) of the 274 sex-biased genes, as judged by comparison of male and female STAT5-KO livers (KOM/KOF comparison; Table S1A). Moreover, sex bias was reduced in the STAT5-KO strain for 80% of the 106 genes that did not meet our cutoff for loss of sex-biased expression. Further supporting our conclusion that STAT5 deficiency impacts the vast majority of sex-biased genes, we found that 81 of the 106 genes that were apparently unresponsive to STAT5-KO actually showed the same overall trend in their response to STAT5 deficiency in male liver as was seen for the 168 significantly responsive sex-biased genes, namely, 43 of 53 male-biased genes decreased in expression, and 38 of 53 female-biased genes increased in expression. Of note, strong female-specific expression was retained in STAT5-KO liver for the GH-regulated transcriptional repressor *Cux2* (43, 44). The absence of an effect on *Cux2* expression may in part explain the incomplete loss of sex-biased gene expression in STAT5-KO livers, given the widespread role of CUX2 in mediating GH-dependent regulation of sex-biased genes in mouse liver (43). Other highly female-biased genes (WTF/WTM >4) that appeared to be truly unresponsive to STAT5 loss (KOM/WTM |fold-change| < 1.2) include *Cyp3a16, Hsd3b1, Ugt2b37* and *Nipal1* (Table S1B).

### STAT5 binding contributes to sex-biased gene expression and STAT5 response

STAT5 binds to liver chromatin in a sex-differential manner nearby a subset of sex-biased genes, as determined by ChIP-seq analysis (12). Here, we examined the set of 98 of 274 sex-biased genes (36%) with a nearby (within 10 kb of gene body) sex-biased STAT5 ChIP-seq peak (Table S1A). Female-biased genes were significantly associated with female-biased STAT5 ChIP-seq peaks, and male-biased genes with male-biased STAT5 ChIP-seq peaks (p<0.0001, Fisher’s Exact test) (**Fig. 1D**, **Fig. 1E**, *left* panels), as expected (12). These positive associations indicate that proximal sex-biased STAT5 binding imparts sex-biased expression, primarily by activation of gene expression. Furthermore, genes down regulated in STAT5-KO male liver were significantly enriched for having proximal male-biased STAT5 ChIP-seq peaks in wild-type liver, consistent with gene down regulation being due to a direct loss of the transcriptional stimulatory effects of STAT5 (**Fig. 1D**, **Fig. 1E**, *middle* panels). In contrast, genes up regulated in STAT5-KO male liver were significantly enriched for proximal female-biased STAT5 ChIP-seq peaks, i.e., STAT5 binding nearby those genes is low in male liver as compared to female liver. This indicates that the STAT5-dependent repression of female-biased genes in wild-type male liver is indirect, e.g., is mediated by a STAT5-dependent repressor whose loss in STAT5-KO liver leads to the observed gene de-repression. The association between gene response to STAT5 deficiency and proximal sex-biased STAT5 ChIP-seq peaks was weaker in female liver (**Fig. 1D**, **Fig. 1E**, *right* panels). STAT5-dependent sex-biased genes that did not show any proximal STAT5 binding (Table S1A) may be regulated by distal STAT5 binding (29).

### Sex-independent genes that respond to STAT5 deficiency

Analysis of sex-independent genes responsive to STAT5-KO gave additional insights into the biological changes occurring in this mouse model. We identified 338 sex-independent genes whose expression was significantly altered in STAT5-KO liver (**Fig. 1A**, Venn regions D, F, G). Many of these genes responded to STAT5 deficiency in the same manner in both sexes, as seen in a heat map displaying 140 of the 338 genes, whose expression was stringently sex-independent in wild-type liver (**Fig. 2A**). Examples include the classic STAT5 target genes *Igf1, Socs2*, and *Onecut1* (HNF6), whose expression was down regulated in livers of both sexes. Pathway analysis identified immunity, cellular response to interferon-β, and peptide antigen binding/MHC as the top enriched pathways for the subset of 71 genes that were induced in common in male and female STAT5-KO livers (**Fig. 2B**, *top*). This contrasts to the strong enrichment of cytochrome P450 and drug and steroid metabolism genes in the set of 155 STAT5-dependent sex-specific genes (**Fig. 2B**, *bottom*). The transcription factor STAT1 was among the genes up regulated in both males and females (**Fig. 2C**), consistent with (28), and may contribute to the observed increases in immune signaling and interferon response gene expression (45). Some residual expression of STAT5b was seen in STAT5-KO liver (**Fig. 2C**), reflecting its continued expression in non-parenchymal cells (42), where the *Alb*-Cre transgene used to excise the STAT5a-TAT5b locus is not active (46).

**Fig. 2 –.**
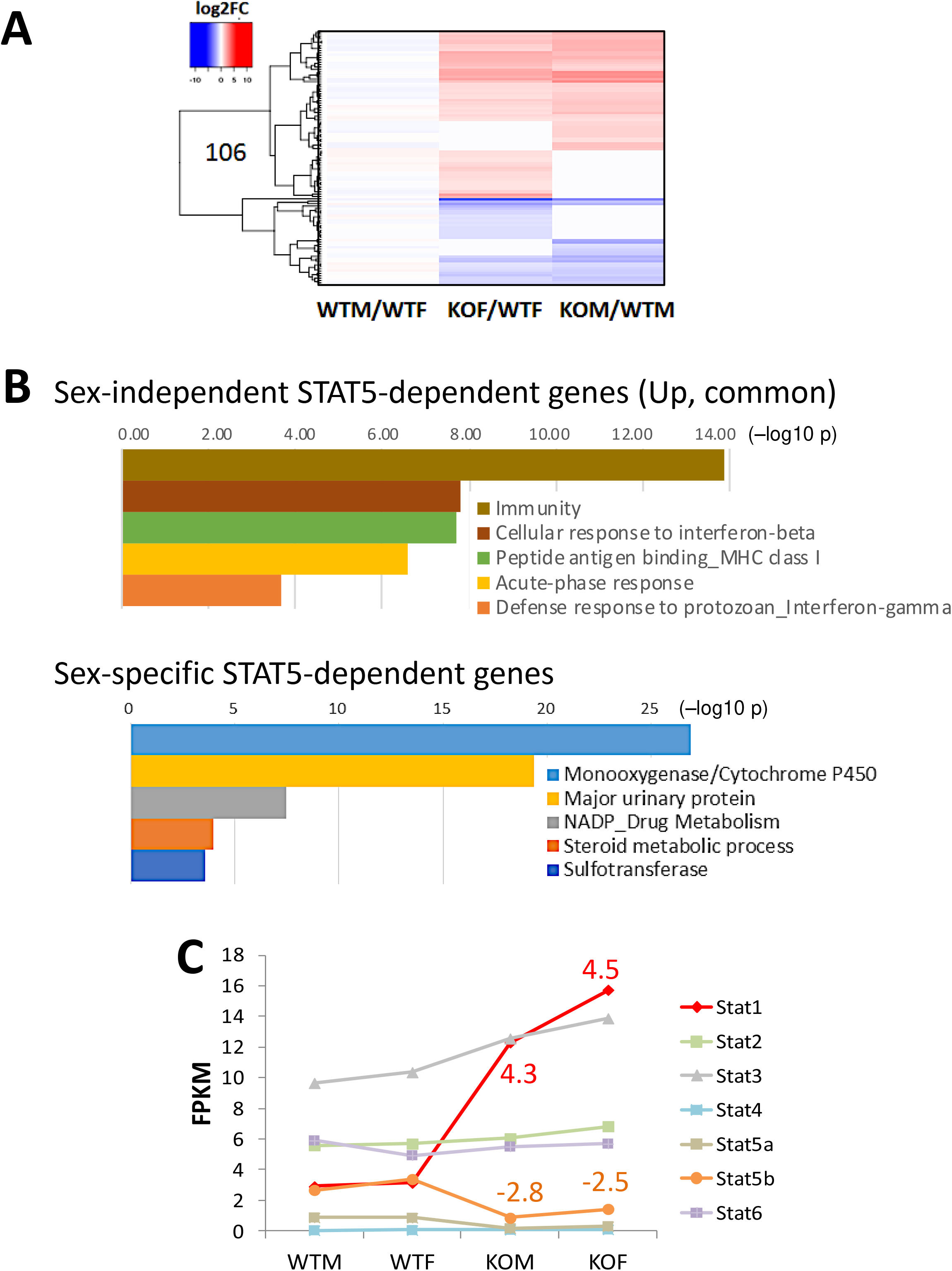
Sex-independent genes responsive to STAT5 knockout (KO). (**A**) Heat map showing impact of STAT5 deficiency on 140 of 6,323 stringent sex-independent genes whose expression was significantly increased (red) or decreased (blue) in either male or female STAT5-KO liver (Table S1A, column S). (**B**) DAVID pathway analysis of 71 genes up regulated in common in STAT5-KO male and female liver (Table S1A, column T) (*top*), and of 155 sex-biased genes responsive to STAT5-KO in male or female mouse liver (Table S1A, column U) (*bottom*). Shown are the top 5 enriched clusters, with bars indicating −log10 p-values (Benjamini-Hochberg corrected) for enrichment. Results are detailed in Table S1C and Table S1D. (**C**) Expression levels of seven STAT family members determined by RNA-seq, expressed as FPKM values. STAT1 was significantly up regulated in both male and female STAT5-KO liver. The incomplete loss of STAT5a and STAT5b is consistent with prior reports (42) and likely reflects STAT5 expressed in non-hepatocytes in mouse liver.

### Impact of STAT5 knockout on DNA methylation in male and female mouse liver

The methylation of state of CpG dinucleotides is a key epigenetic mark regulating chromatin structure and gene expression. Given the major impact of liver STAT5 deficiency on sex-biased gene expression, shown above, we carried out reduced representation bisulfite sequencing (RRBS) to capture and analyze the methylation status of genomic regions with high CpG content in wild-type and STAT5-KO mouse liver. We used the same set of livers as in the RNA-seq analysis, but analyzed a larger number of individual livers (biological replicates; N=5) to increase statistical power. We identified genomic regions that show significantly differential CpG methylation (DMRs, differentially methylated regions), either between male and female wild-type mouse liver (sex-biased DMRs), or between STAT5-KO and wild-type mouse liver (STAT5-dependent DMRs) (Table S2).

Each DMR covers a 100 bp genomic region (‘tile’), which may be either hypermethylated or hypomethylated in the treatment sample compared to its control (e.g., male compared to female, or STAT5-KO compared to wild-type). The extent of differential methylation was quantified as a differential methylation (diff.meth) value, corresponding to the difference in percent methylation between the samples being compared (**Fig. 3A**). Hypermethylation of the treatment relative to control is indicated by a positive diff.meth value; and hypomethylation, where treatment is demethylated relative to control, is indicated by a negative diff.meth value. DMRs with diff.meth between |15%| and |25%| at FDR < 0.05 were considered significant at standard stringency; diff.meth > |25%| at FDR < 0.05 identified robust DMRs, as shown in **Fig. 3A** for each of four pair-wise comparisons. DMRs sensitive to the loss of STAT5 were more frequently associated with hypermethylation than hypomethylation when compared to wild-type liver, most notably in males (Hyper/Hypo methylation ratio = 2.14 for KOM/WTM vs. 0.9-1.01 for WTM/WTF and KOM/KOF). The trend of increased hypermethylation with STAT5 deletion was also apparent by boxplot analysis (**Fig. 3B**). Furthermore, unsupervised (hierarchical) clustering of raw methylation values for all robust DMRs identified in the KOM/WTM comparison correctly separated all of the male from female wild-type individuals, and all but one of the STAT5-KO from wild-type individuals (**Fig. 3C**, heat map).

**Fig. 3 –.**
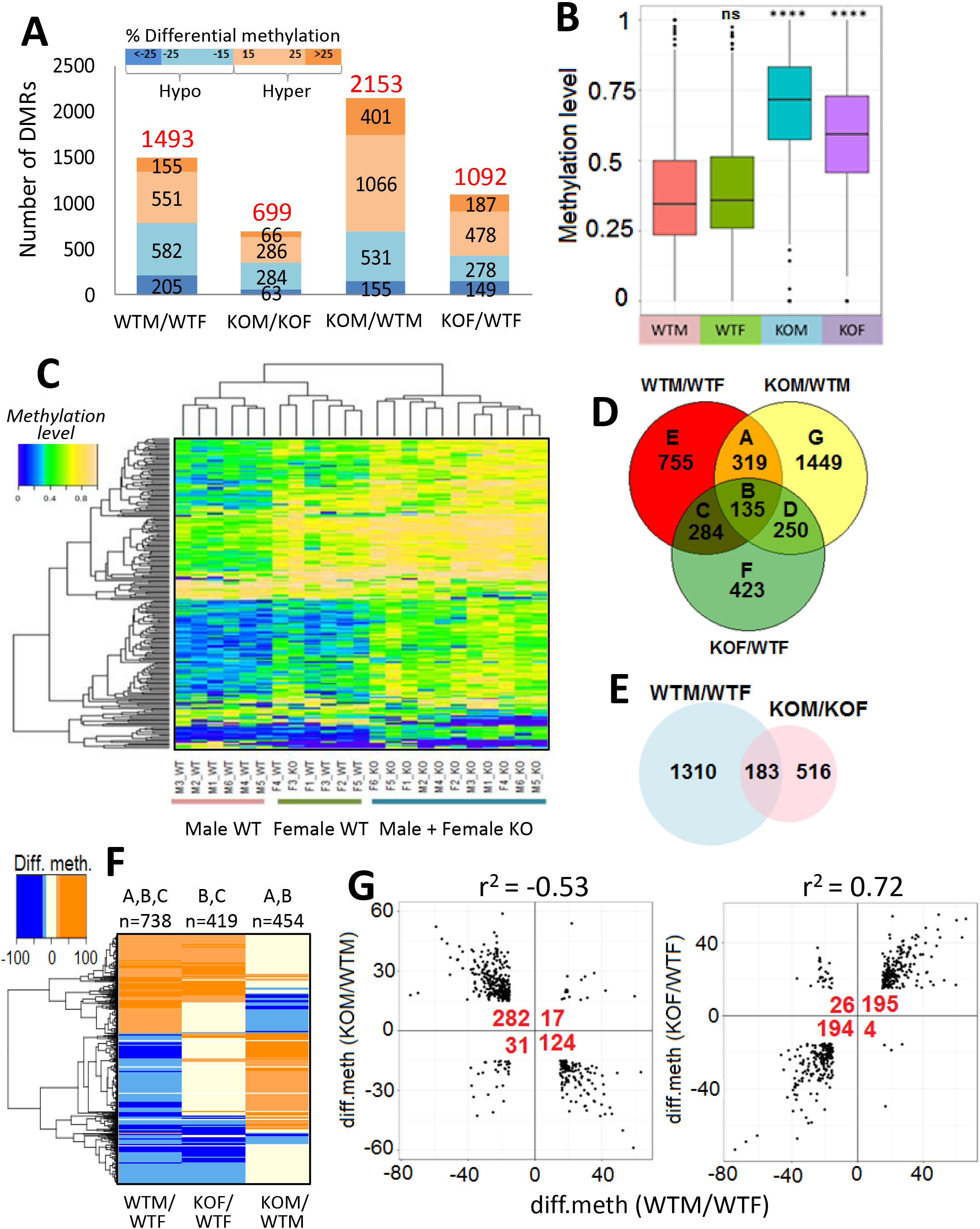
DNA methylation changes in STAT5-KO compared to wild-type mouse liver. (**A**) Shown are the numbers of all DMRs (based on both tile statistic and individual CpG-based DMRs, i.e., the combined set in Table S2F) that were hypermethylated (orange) or hypomethylated (blue) at standard (|15-25|%) or robust (>|25|%) thresholds for differential methylation in each of the four indicated comparisons. (**B**) Box plots of the raw methylation levels of each of the four indicated groups, for the set of 383 tile-based DMRs that exhibited > 25% diff.meth between STAT5-KO and wild-type male liver, showing that STAT5 loss largely results in hypermethylation. ****, p < 0.0005 for Student t-test comparison of STAT5-KO to the sex-matched wild-type group. Analysis is based on n=5-6 individual livers per group, as shown in C. (**C**) Heat map showing raw methylation levels for the 383 DMRs analyzed in B. Shown are datasets for 23 individual livers, subjected to hierarchical clustering. Color bar: raw methylation levels range from 0% methylation (blue) to 100% methylation (yellow). (**D**) Venn diagram showing the overlap of sex-biased DMRs in wild-type liver compared to STAT5-KO male and female liver, representing a total of 3,615 DMRs (>15% diff. meth). The number of DMRs in each comparison (labeled A through G) is indicated. (**E**) Venn diagram overlap of sex-biased DMRs in wild-type compared to STAT5-KO mouse liver. Analyses in D and E are based on both tile statistic and individual CpG-based DMRs. (**F**) Heat map showing hypermethylated DMRs (orange) and hypomethylated DMRs (blue) from Venn diagram comparisons A, B and C in panel D. (**G**) Scatter plots showing the relationship between sex-specificity and STAT5 responsiveness of DMRs. Left graph: 454 sex-biased DMRs, from panels D and E, that are responsive to loss of STAT5 in male liver. Of the sex-biased DMRs that are hypomethylated in wild-type male compared to female liver, 282 show increased methylation and 31 show increased demethylation in STAT5-KO male liver, and those that are hypermethylated in wild-type male compared to female liver, 124 become demethylated and 17 show increased methylation in STAT5-KO male liver. Right graph: 419 sex-biased DMRs, from panels D and E, that are responsive to STAT5 loss in female liver. STAT5 deficiency results in methylation and demethylation of these DMRs with response patterns opposite to those shown in left graph. Fisher’s exact test for DMR responses between the four quadrants: p<2.2E-16 for both graphs. Linear regression analysis of |diff. meth| values showed significant correlations between magnitude of sex-specificity and response to STAT5-KO (r^2^ values).

Of the 1493 sex-differential DMRs seen in wild-type liver, 738 (49%) showed significant differential methylation between STAT5-KO and wild-type livers, i.e., are STAT5-responsive DMRs, in either males or females (**Fig. 3D**; sections A+B+C; **Fig. 3F**). Moreover, STAT5 deficiency caused a loss of significant sex-differences in CpG methylation at 1310 (88%) of the 1493 sex-differential DMRs (WTM/WTF vs KOM/KOF comparison; **Fig. 3E**). In male STAT5-KO liver, DMRs with a significantly lower methylation level in wild-type male compared to wild-type female liver (i.e., male-hypomethylated DMRs) primarily became hypermethylated, while many male-hypermethylated DMRs became hypomethylated (**Fig. 3F** and **Table 2**). In contrast, in female STAT5-KO liver, many female-hypermethylated DMRs (i.e., male-hypomethylated DMRs) became hypomethylated, and female-hypomethylated DMRs became hypermethylated. **Fig. 3F** shows that STAT5-KO induces a loss of sex-differential methylation at male-hypomethylated sites by increasing methylation in male liver or by decreasing methylation in female liver. In the case of female-hypomethylated sites, STAT5-KO induces a loss of sex-differential methylation by increasing methylation in female liver or by decreasing methylation in male liver. Quantitative scatter plots illustrate this significant reversal of sex-biased differential DNA methylation in the absence of liver STAT5 (**Fig. 3G**). Further, the extent of methylation differences between sexes correlated significantly with the level of DNA methylation changes in the STAT5-KO livers (**Fig. 3G**). Thus, liver STAT5 is required for sex-specific methylation and demethylation at several hundred genomic regions.

**Table 2.**
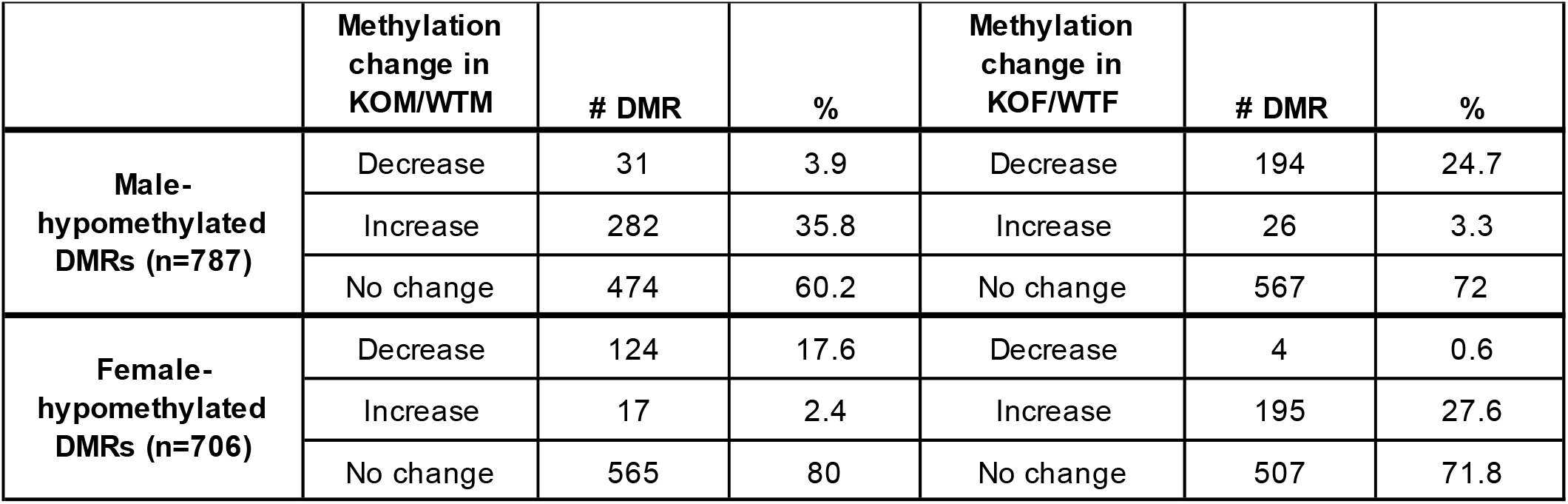
Differential methylated regions in hepatocyte-specific STAT5-KO liver. 1493 sex-biased DMRs (**Fig. 3D**) were classified into DMRs that were more hypomethylated in male liver (787) or more hypomethylated in female liver (706). These DMRs were further sorted by their direction of change in STAT5-deficient compared to wild-type liver, as indicated, in both male liver (KOM/WTM) and in female liver (KOF/WTF).

|  | Methylation<br>change in<br>KOM/WTM | # DMR | % | Methylation<br>change in<br>KOF/WTF | # DMR | % |
| --- | --- | --- | --- | --- | --- | --- |
| **Male-**<br>**hypomethylated**<br>**DMRs (n=787)** | Decrease | 31 | 3.9 | Decrease | 194 | 24.7 |
|  | Increase | 282 | 35.8 | Increase | 26 | 3.3 |
|  | No change | 474 | 60.2 | No change | 567 | 72 |
| **Female-**<br>**hypomethylated**<br>**DMRs (n=706)** | Decrease | 124 | 17.6 | Decrease | 4 | 0.6 |
|  | Increase | 17 | 2.4 | Increase | 195 | 27.6 |
|  | No change | 565 | 80 | No change | 507 | 71.8 |

### Few DMRs are in promoter regions of sex-biased, STAT5-dependent genes

CpG islands, found at many gene promoters, generally exhibit low methylation levels and are correlated with active transcription (47). Further, hypermethylation of CpG islands is associated with stable silencing of genes (48). To determine if promoter CpG methylation contributes to the sex-biased and STAT5-dependent gene expression changes we observed, we examined DMRs proximal (5 kb upstream or downstream) to the TSS of sex-biased, STAT5-responsive genes. Very few such sex-biased genes have a proximal sex-biased DMR (**Fig. 4A**). Further, only two STAT5-dependent, male-biased genes, *Cyp7b1* and *Cspg5*, showed significant promoter DNA methylation changes in STAT5-KO male liver that matched the general trends described above (**Fig. 4B**, **Fig. 4C**, Table S3A). For these two genes, DNA methylation in the promoter region was significantly lower in wild-type male than female liver, and was increased in STAT5-KO male liver in association with the decrease in gene expression. Thus, the large majority of sex-biased, STAT5-dependent genes do not display sex differences in promoter region DNA methylation. We also identified examples of sex-independent STAT5-KO responsive genes that showed STAT5-dependent changes in promoter region DNA methylation in a direction consistent with the change in their expression in STAT5-deficient male and/or female liver (Table S3B).

**Fig. 4. –.**
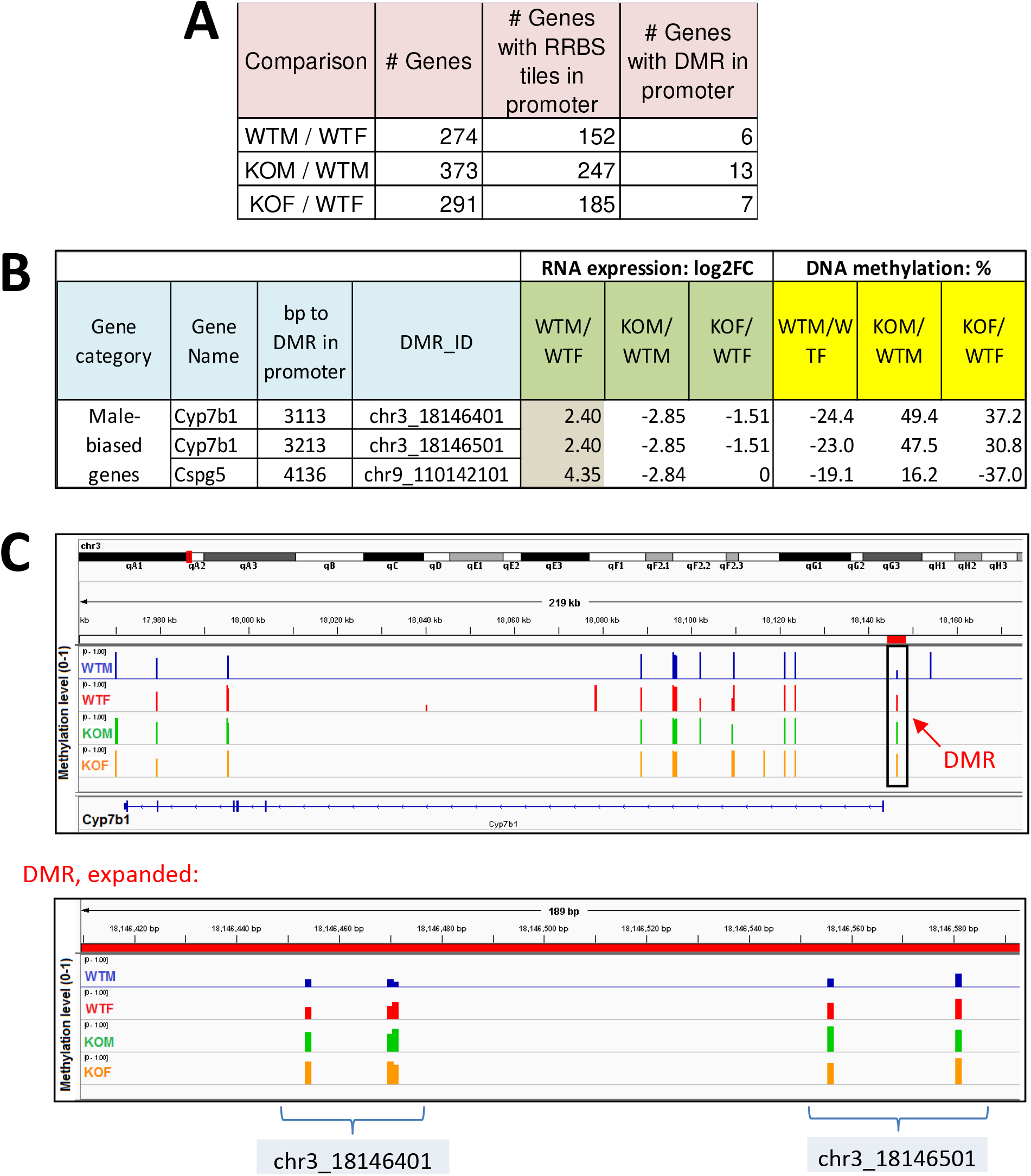
Few DMRs are at promoters of sex-biased or STAT5-dependent genes. (**A**) Shown are the number of significant differentially expressed genes identified in each comparison (as in Fig. 1A), for genes with an RRBS-covered tile or with a significant DMR tile within 5 kb upstream of gene transcription start site (promoter region). (**B**) Differential expression and differential methylation data for promoter DMR/gene pairs that are consistently male-biased at both the gene expression level and the promoter DMR level. FC, fold-change. (**C**) Browser screen shot of genomic region encompassing *Cyp7b1*, with the methylation level of individual RRBS tiles shown for male and female wild-type and for male and female STAT5-KO liver. Only the 3’-most RRBS tiles (boxed) correspond to DMRs. The latter region is expanded at the bottom and encompasses two adjacent DMRs, each containing 4 CpGs, whose percent methylation is indicated along the y-axis for each of the four conditions. The extent of differential methylation for these CpGs is shown in B.

### Chromatin states and chromatin accessibility at DMRs

To determine the chromatin context of STAT5-dependent DNA methylation, we mapped each set of DMRs to 14 ChromHMM-derived chromatin states (**Fig. 5A**) previously learned for male and female mouse liver using genome-wide datasets for open chromatin regions (DHS, DNase-I hypersensitive sites) and histone marks (29). A substantial fraction of the >1.3 million CpGs captured by our RRBS samples were within a promoter state, reflecting the preferential capture of promoter CpG island sequences by the RRBS protocol. Promoter states show low CpG methylation overall, while enhancer states show a large range of CpG methylation levels (Fig. S2). CpGs in sex-biased and STAT5-responsive DMRs were significantly depleted of promoter states (blue) and showed various levels of enrichment for enhancer states (green) (**Fig. 5B**, **Fig. 5C**). Frequency profiles of the 14 chromatin states in a 4-kb window surrounding each DMR revealed specific enrichment of enhancer state 6 for DMRs that were hypermethylated, but not hypomethylated, in the absence of STAT5 (KOM/WTM and KOF/WTF comparisons) (**Fig. 5D**). A modest enrichment of enhancer state 6 was also seen in DMRs hypomethylated in wild-type male relative to wild-type female liver. Enhancer state 6 represents active enhancers characterized by high probabilities of H3K4me1 and H3K27ac histone marks and presence of a DHS (**Fig. 5A**). Thus, loss of STAT5 leads to a preferential hypermethylation of genomic regions in an active enhancer state.

**Fig. 5 –.**
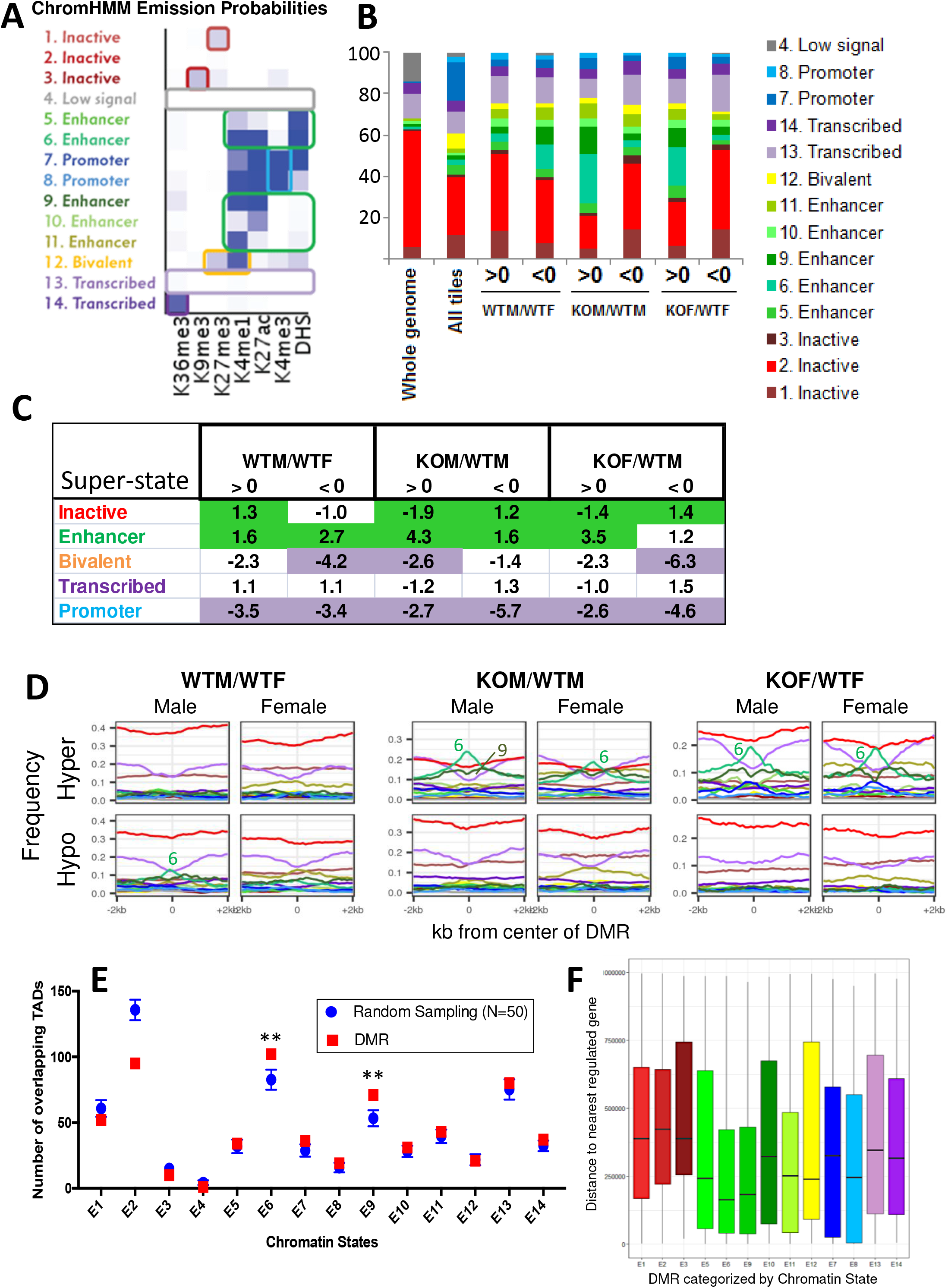
Chromatin states at DMRs responsive to STAT5-KO in liver. (**A**) ChromHMM emission probabilities for the 14 chromatin states discovered in male and female mouse liver, from (29). Each state is colored and given a descriptive name (*left*) based on the frequency of each of six chromatin mark and presence of DHS, as indicated at the bottom. (**B**) Cumulative distribution of chromatin states for the set of 232,882 tiles captured by the 23 RRBS samples used in this study, and for the subsets of tiles identified as DMRs in the WTM/WTF, KOM/WTM, and KOF/WTF comparisons. The first bar shows that the whole mouse genome is mostly covered by chromatin state 2, an inactive state. All Tiles indicates the chromatin state distribution of the full set of 232,882 tiles, which is most strongly enriched for promoter states (blue) and poised states (yellow) when compared to the whole genome. DMRs identified in each of the 3 pairwise comparisons are marked >0 (hypermethylation) or <0 (hypomethylation), and show a relative depletion of promoter states and variable increases in the proportions in enhancer states (various shades of green). For example, hypermethylated DMRs in the KOM/WTM and KOF/WTF datasets showed an increased proportion of enhancer states. (**C**) Enrichment scores for inactive, enhancer, bivalent, transcribed and promoter chromatin states in each DMR group relative to the full set of tiles captured by RRBS. To calculate enrichment scores, the 14 ChromHMM chromatin states were grouped into five super states, designated as inactive (states 1, 2, 3), enhancer (states 5, 6, 8, 9, 10), bivalent (state 12), transcribed (states 13, 14) and promoter (states 7, 8). Significant enrichment is shown in green and significant depletion in purple (p-value cutoff: 1E-5, two proportion z-test). (**D**) Frequency of occurrence of each of the 14 chromatin states in male and female liver, colored as indicated in B, was calculated in 100 bp windows extending 2 kb upstream and 2 kb downstream of the hypermethylated and hypomethylated DMRs in each comparison. Chromatin state 6 showed a prominent peak in hypermethylated DMRs for the KOM/WTM and KOF/WTF comparisons, in both male and female liver, and to a less extent, in hypomethylated DMRs of the WTM/WTF comparison in the male liver, as marked. (**E**) Genes that are sex-biased or STAT5-responsive show a significant enrichment for being in the same TAD as a sex-biased and/or STAT5-responsive DMR. **, Z-score >2.5. Inactive chromatin state E2 shows fewer overlapping TADs than a random sampling of CpG regions (also see Table S4C). (**F**) Box plots showing distribution of distances to the nearest sex-biased and/or STAT5-responsive gene TSS for sex-biased and/or STAT5-responsive DMRs grouped by their chromatin state, as shown along the x-axis. See Table S4 for datasets used in these analyses.

### TAD-based analysis of association of sex-biased and STAT5-dependent genes with distal DMRs

Next, we investigated whether the enhancer state-associated DMRs can be linked to STAT5-dependent transcriptional regulation. As it is difficult to directly link a given enhancer to its specific gene target(s), we evaluated such interactions at the level of topologically associating domains (TADs), which are megabase-scale genomic regions with high propensity for intra-domain contact in three-dimensional chromatin organization (49). Using TAD boundaries defined for mouse liver (50, 51), we investigated whether genes that are sex-biased or STAT5-responsive are more likely to be in the same TAD as a sex-biased or STAT5-responsive DMR, when compared to sets of random CpG regions. DMRs, grouped by chromatin state (**Fig. 5B**), were mapped to TADs, and their overlap with the set of 437 TADs containing at least one sex-biased or STAT5-responsive gene was then determined (Table S4). The statistical significance of TAD overlap was determined for each chromatin state by comparison to TAD overlap for a random sampling, performed 50 times, of comparably sized sets of 100 bp RRBS-captured CpG regions that did not show significant differential methylation. DMRs in chromatin states 6 and 9, but not DMRs in other chromatin states, showed significantly higher co-occupancy with TADs containing sex-biased or STAT5-responsive genes than did random CpG regions (z-score: 2.54 and 2.98; p-value: 0.011 and 0.004, for chromatin states 6 and 9, respectively) (**Fig. 5E**, Table S4C). Chromatin states 6 and 9 are enhancer states with high levels of histone H3K4me1 and histone H3K27ac marks, either with (state 6) or without (state 9) a DNase hypersensitive site (DHS) (**Fig. 5A**), with the DHS-free enhancer state 9 showing much higher overall CpG methylation than enhancer state 6 (Fig. S2). These enrichments support the proposal that the regulation of CpG methylation at these gene-distal enhancers plays a regulatory role for the sex-biased and STAT5-responsive genes within these TADs. Consistent with this proposal, the sex-biased, STAT5-responsive DMRs in chromatin states 6 and 9 were closer in median distance to sex-biased, STAT5-responsive genes than DMRs in other chromatin states (**Fig. 5F**).

### Association of DMRs with nearby open chromatin regions

Many DMRs were located nearby at least one liver DHS (Fig. S3). These DMR-proximal DHS include 308 male-biased DHS and 66 female-biased DHS identified previously (52) (Table S5). Gene set enrichment analysis showed that these 374 sex-biased DHS were non-randomly distributed: the 308 male-biased DHS were associated with DMRs that were hypomethylated in wild-type male liver (normalized enrichment score, NES: −3.18, FDR=0), and the 66 female-biased DHS were associated with DMRs hypermethylated in wild-type male liver (and consequently, hypomethylated in wild-type female liver) (NES=1.57, FDR=0.007) (**Fig. 6**, *left* panels). Thus, sex-biased CpG demethylation is associated with sex-biased chromatin opening in wild-type livers of both sexes. Moreover, in male liver, loss of STAT5 resulted in significant hypermethylation in genomic regions with a male-biased DHS, but not in region with female-biased DHS (*middle* panels), whereas the absence of STAT5 in female liver resulted in hypermethylation at sites nearby both male and female DHS (*right* panels). Thus, sex-biased chromatin opening is associated with sex-biased CpG demethylation, and loss of liver STAT5 enables hypermethylation of nearby sex-biased DHS, consistent with the enrichment of the DHS-containing enhancer state 6 seen in **Fig. 5D**.

**Fig. 6 –.**
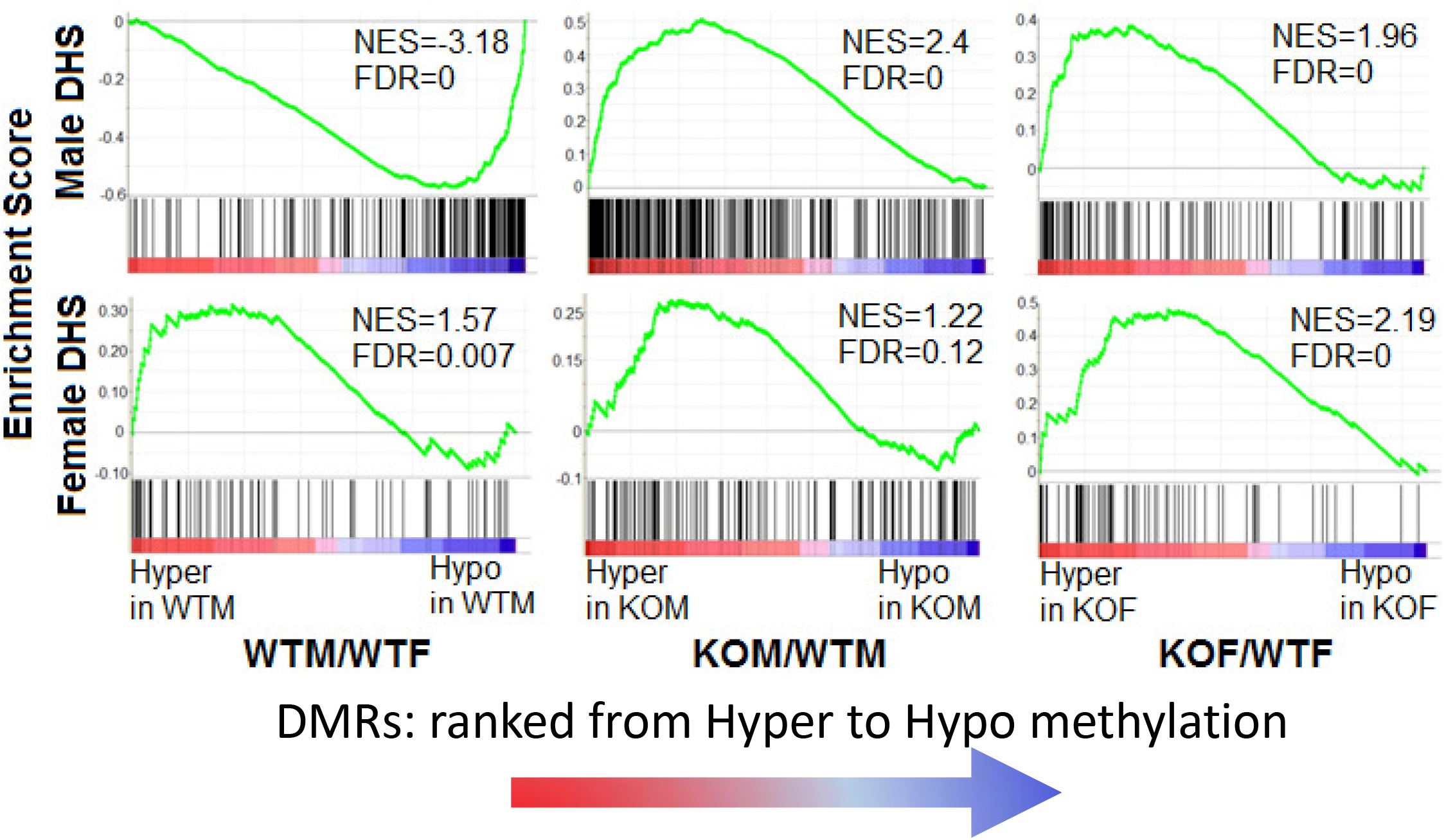
GSEA analysis of STAT5-responsive DMRs nearby sex-biased DHS. DMRs in the three comparisons indicated at the bottom were ranked from hypermethylation (red) to hypomethylation (blue). GSEA was used to determine whether sex-biased DHS (Table S5) are preferentially found within 5 kb of the hypermethylated or the hypomethylated end of the ranked DMR spectrum. NES: normalized enrichment score; FDR < 0.05 was considered significant. For example, DMRs that were hypomethylated in male liver were enriched for male-biased DHS (NES: −3.18, FDR=0), while DMRs hypomethylated in male liver (and thus hypomethylated in female liver) were enriched for female-biased DHS (NES=1.57; FDR=0.007).

### STAT5 binding nearby DMRs

Given our finding that the loss of STAT5 in male liver increases methylation at male-biased DHS regions associated with hypomethylated DMRs, we scanned each set of DMRs (WTM/WTF; KOM/WTM; KOF/WTF) for STAT5 motifs. We found strong enrichment of the STAT5 motif at hypermethylated DMRs in STAT5-KO male liver, and to a lesser extent in STAT5-KO female liver, but not at the corresponding hypomethylated DMRs, consistent with hypermethylation occurring at or nearby STAT5 binding sites (**Fig. 7A**). Next, we examined whether STAT5 ChIP-seq binding sites identified in wild-type male and female liver (12) are enriched in the set of DMRs hypermethylated in STAT5-KO liver. Overall, 59% of all liver STAT5 ChIP-seq peaks were nearby one or more of the CpG sites covered by our RRBS dataset (**Fig. 7B**). Cumulative frequency plots of distance to the nearest STAT5 binding site (**Fig. 7C**) showed that a much higher fraction of DMRs that were hypermethylated in STAT5-KO liver were directly at (y-intercept values; solid lines) or nearby a STAT5 binding site as compared to DMRs that were hypomethylated (dashed lines), consistent with the STAT5 motif analysis. When considering STAT5 binding sites within a few kb of a DMR, male-biased STAT5 binding sites showed greater enrichment for DMRs hypermethylated by STAT5-KO in male liver than those hypermethylated in STAT5-KO female liver (**Fig. 7C**, *middle*), while female-biased STAT5 binding was most enriched in hypermethylated DMRs found in STAT5-KO female liver (**Fig. 7C**, *bottom*). Further, male-biased STAT5 binding sites (and to a lesser extent sex-independent STAT5 sites) were preferentially enriched at DMRs hypomethylated in male liver (*middle*, purple dashed line). We conclude that STAT5 binding sites are preferentially hypomethylated in wild-type male, but not female liver, and that the loss of STAT5 increases CpG methylation proximal to STAT5 binding in both male and female liver.

**Fig. 7 –.**
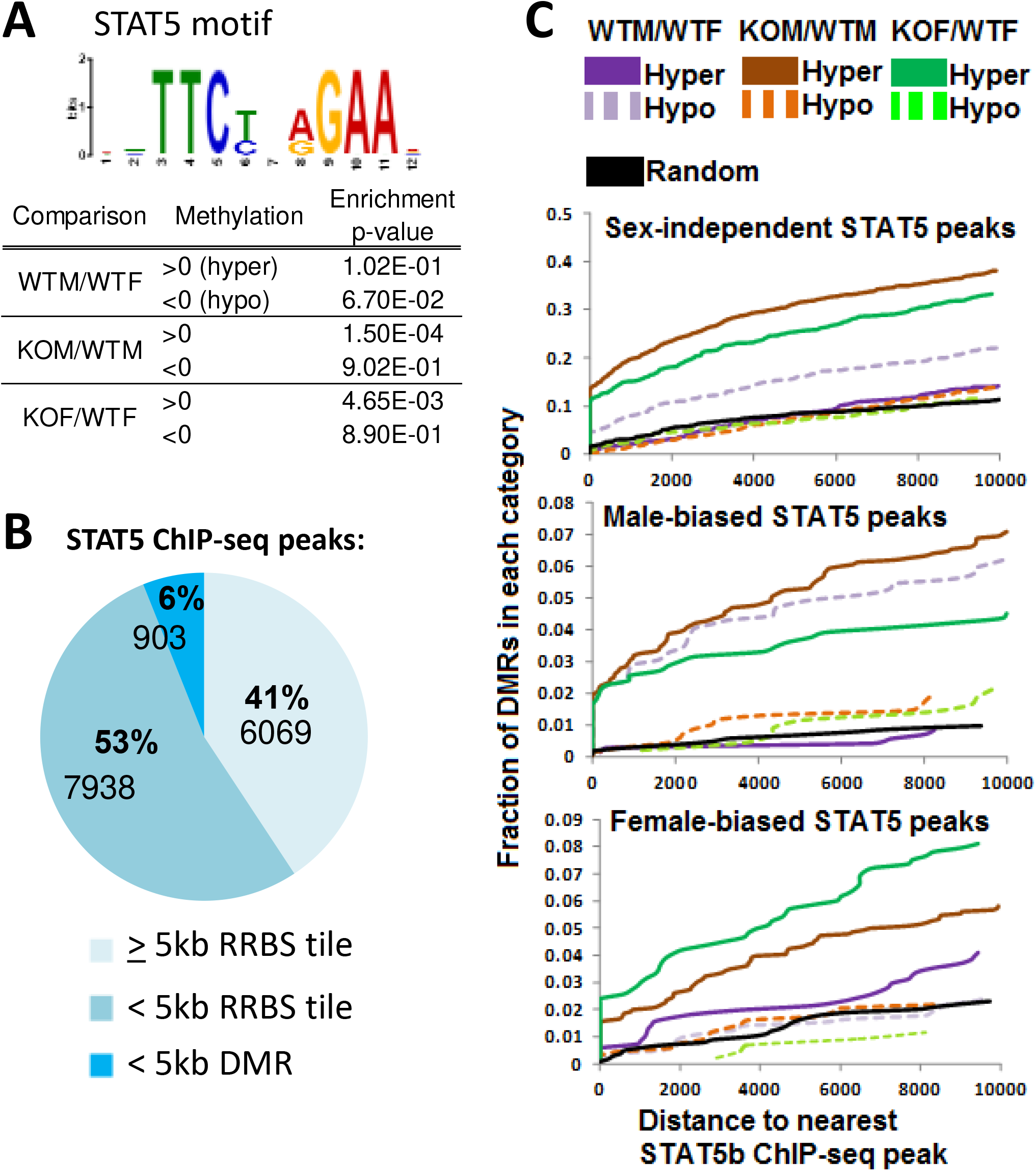
STAT5 motifs and ChIP-seq peaks at DMRs. (**A**) Enrichment of STAT5 motifs at DMRs determined using the MEME Analysis of Motif Enrichment tool. (**B**) Categorization of 14,910 STAT5 ChIP-seq sites in mouse liver (12) based on whether they are within 5 kb of one of the 232,882 RRBS tiles captured in this study, or whether they are within 5 kb of one of the RRBS that is also a DMR. Here, DMR refers to 3,615 regions that showed significant differential methylation in one or more of these three comparisons: WTM/WTF, KOM/WTM, and KOF/WTF. (**C**) Cumulative frequency plots of hypermethylated and hypomethylated sex-biased and STAT5-reponsive DMRs and their distances to the nearest STAT5 binding site in each set of sites identified in B (Table S2F, last columns).

### Motifs enriched at hypermethylated DMRs

Next, we used *de novo* motif discovery, implemented using Homer (53), to determine whether motifs for other transcription factors are enriched in the sets of sex- or STAT5-dependent hypermethylated and hypomethylated DMRs described above. The largest number of significantly enriched motifs (n = 47, q-value <0.05) was discovered in the set of hypermethylated DMRs from the KOM/WTM comparison, followed by 13 motifs for hypermethylated DMRs from the KOF/WTF comparison (**Fig. 8A**). Analysis of the overlaps of enriched motifs between datasets revealed that a majority of the hypermethylated KOF/WTF motifs overlapped the set of hypermethylated KOM/WTM motifs (**Fig. 8B**). The top motifs discovered are shown in **Fig. 8C**. Notably, motifs for STAT5, HNF6 and CUX2, all previously implicated in regulation of liver sex differences (29, 43, 44), were unique to the sets of hypermethylated DMRs common to the KOM/WTM and KOF/WTF comparisons. Thus, loss of STAT5 resulted in hypermethylation of regions enriched in DNA motifs for STAT5 and its co-regulating transcription factors. These regions are enriched for enhancer histone marks and chromatin states (**Fig. 5**), DHS (**Fig. 6**), and STAT5 binding (**Fig. 7**) in wild-type liver, and their presumed loss of activity upon hypermethylation (54, 55) is expected to contribute to the loss of sex-biased expression of their associated genes.

**Fig. 8 –.**
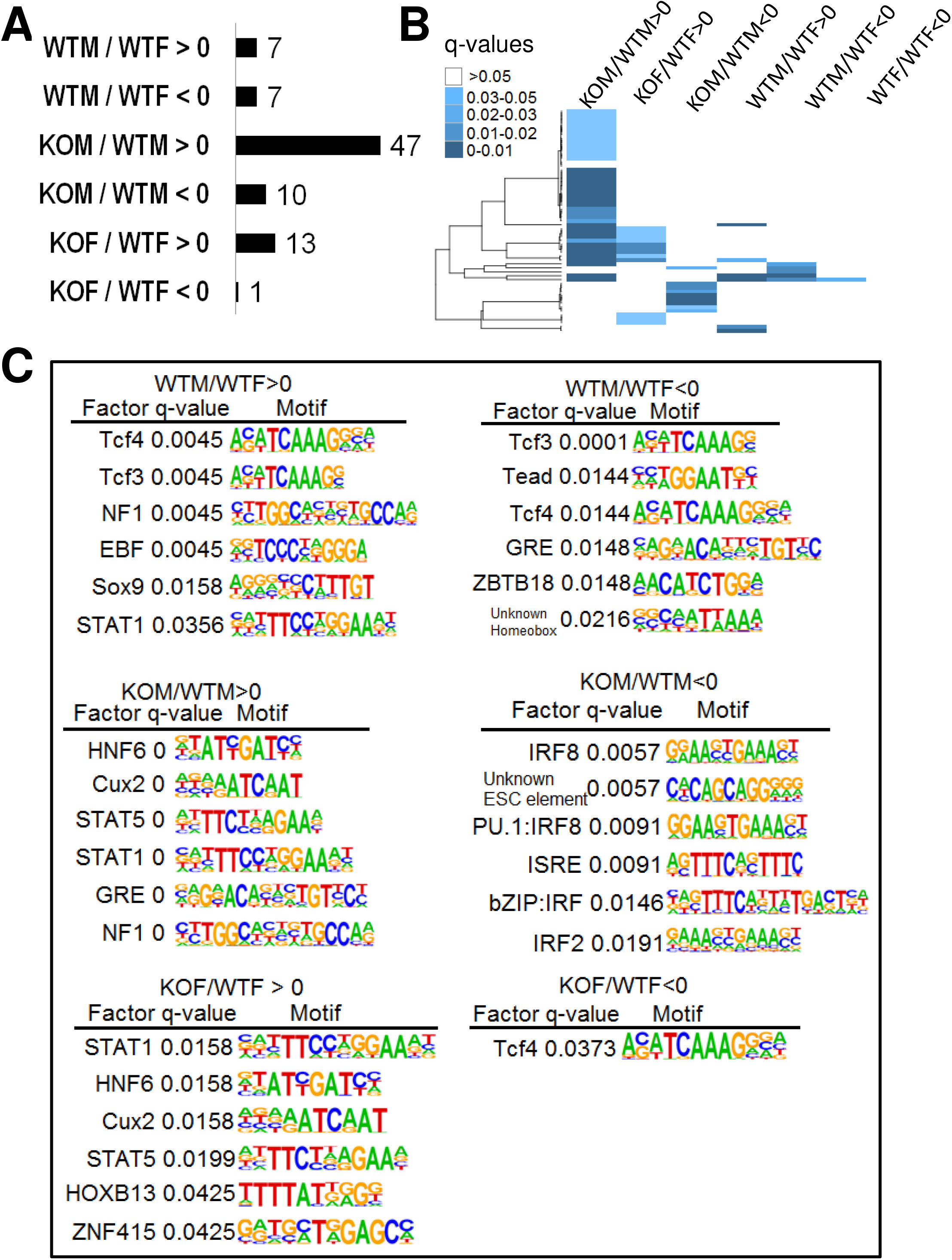
Enriched DNA motifs at DMRs. (**A**) Number of enriched DNA motifs determined by Homer that passed the significance filter of q-value >0.05. (**B**) Heat map showing the clustering of DNA motifs numbered in A, indicating the extent of overlap between the 6 indicated sets of motifs (top). (**C**) Motif name, q-value and sequence logo for the top six *de novo*-discovered DNA motifs (lowest q-value) for the indicated 6 sets of hypermethylated and hypomethylated DMRs. See Table S6 for a complete listing of significant motifs.

## Discussion

We investigated the relationship between sex-biased DNA methylation and sex-biased liver gene expression in mice with a hepatocyte-specific deletion of STAT5, which plays a central role in GH regulation of adult liver sexual differentiation. Loss of STAT5 in hepatocytes led to a substantial, albeit incomplete loss of sex-biased liver gene expression, with STAT5 inducing male-biased genes in male liver via a positive, STAT5-binding-dependent regulatory mechanism, and with STAT5 repressing female-biased genes in male liver by an indirect regulatory mechanism. Liver STAT5 deficiency also resulted in widespread changes in DNA methylation, abolishing the sex bias in DNA methylation at 88% of 1,493 sex-biased DMRs. Hypermethylated regions identified in male and female STAT5-KO liver were enriched for open chromatin regions (DHS), enhancer chromatin states, and STAT5 ChIP-seq peaks in wild-type liver, and encompassed transcription factor motifs for STAT5 and other transcription factors previously implicated as regulators of liver sex bias. Thus, many DMRs that become hypermethylated in STAT5-KO liver represent STAT5-bound enhancers that require STAT5 to maintain a low DNA methylation state. As DNA methylation is often associated with condensed chromatin that is inaccessible to transcription factor binding (56), our findings suggest that the loss of STAT5 leads to silencing of these regulatory elements via CpG methylation. Proximal promoter regions of STAT5-responsive genes showed very few significant changes in DNA methylation in STAT5-KO liver, and correspondingly, promoter states were significantly under-represented, especially in the sets of DMRs that become hypomethylated in the absence of STAT5. The later DMRs may represent latent regulatory elements that become activated in the absence of STAT5, e.g., due to the up regulation of STAT1 (**Fig. 2**), or by other compensatory mechanisms that have yet to be identified.

The present findings in hepatocyte-specific STAT5a/STAT5b-KO liver differ somewhat from prior findings in whole body STAT5b-KO liver based on microarray analysis (10), where liver sex bias was abolished for a larger fraction of sex-biased genes (78% vs. 61%). However, a full 80% of genes that apparently retained sex-biased expression in the present hepatocyte-specific STAT5-KO model actually showed a decrease in the magnitude of their sex bias in the absence of STAT5. Furthermore, and consistent with our prior work (10), the impact of hepatocyte-specific STAT5 loss was less pronounced in female liver, where fewer genes were dysregulated, and where there was a weaker correlation between sex specificity and STAT5 response and a weaker association with sex-biased proximal STAT5 binding. Nevertheless, hepatocyte-specific STAT5 deficiency did impact sex-biased gene expression in female liver, consistent with the female-biased binding of STAT5 to liver chromatin seen by ChIP-seq (12) and with the finding that STAT5a is specifically required for expression of 15% of female-specific genes in female liver (57).

One factor contributing to the more extensive loss of sex-biased gene expression in the whole body STAT5b-KO mouse model (10) may be the loss of STAT5 throughout prenatal and postnatal development, as well as its absence in all liver cell types, including non-parenchymal cells. In contrast, in the present, hepatocyte-specific STAT5 knockout mouse model, STAT5 deletion was achieved by activation of a Cre transgene under the control of the *Alb* gene promoter (28), which does not occur in liver non-parenchymal cells, and consequently, there is significant residual expression of STAT5 (42). Another factor contributing to the less extensive loss of sex-biased gene expression in the hepatocyte-specific STAT5-KO mouse model may be CUX2, a female-specific transcriptional repressor that is rapidly induced by continuous GH infusion in male liver (21) and is de-repressed in whole body STAT5b-KO male liver (58), but not in hepatocyte-specific STAT5-KO mouse liver, as shown here. The induction of CUX2 seen in whole body STAT5b-KO male liver is likely due to its direct, STAT5-independent induction by the persistent elevation of circulating GH found in that mouse model (42), which would enable CUX2 to contribute to a more complete feminization of male mouse liver through its widespread repression of male-biased genes (43). Other strongly female-biased genes that were not induced in hepatocyte STAT5-deficient male liver include the drug metabolizing genes *Cyp3a16, Hsd3b1* and *Ugt2b37*. Given that these genes, as well as *Cux2*, are strongly induced in livers of male mice treated with GH as a continuous infusion (female plasma profile) (21), it seems that their sex-biased expression is regulated by a GH-dependent but STAT5-independent signaling pathway.

CpG methylation at a specific promoter-proximal CpG dinucleotide has been implicated in the regulation of two sex-biased liver genes, *Cyp2d9* and *C4a/Slp* (38, 59, 60). Our studies identified differential methylation proximal to *C4a*/Slp (Table S2F); however, our RRBS analysis did not identify CpGs nearby *Cyp2d9*, which could be due to insufficient CpG density in that region. While many sex-biased, STAT5-responsive gene promoters contain nearby CpGs captured by our RRBS analysis, very few differentially methylated CpG tiles were found at such promoters. Rather, our findings indicate that STAT5-dependent sex-biased genes are frequently regulated by STAT5-regulated demethylation associated with the activation of distal enhancers. Thus, we observed a high correlation between the sex-bias and STAT5 responsiveness of CpG methylation changes at sex-biased DMRs in both male and female liver. Loss of STAT5 preferentially increased CpG hypermethylation, with the sites that are hypermethylated in the absence of STAT5 being more strongly enriched for enhancer chromatin states, presence of open chromatin (DHS), and STAT5 binding as compared to hypomethylated sites. This indicates that de-methylation at regulatory elements is dependent on STAT5, and that loss of STAT5 leads to retention of DNA methylation at genomic loci that would otherwise be active, demethylated enhancers that bind STAT5 and undergo trans-activation. This regulatory mechanism encompasses many distal enhancers, as indicated by the significant enrichment of DMRs in enhancer chromatin states for co-localization with sex-biased and/or STAT5-responsive genes within their TAD boundaries. Whether DNA demethylation plays a direct, instructive role in promoting STAT5 binding, or is a passive consequence of binding of STAT5 or associated transcription factors is unknown.

The widespread effects of STAT5a/STAT5b-KO on liver sex differences in gene expression and DNA methylation seen here can primarily be attributed to the loss of STAT5b, which is GH-responsive and is the predominant liver STAT5 form. STAT5a is also GH responsive, but is expressed in mouse liver at a much lower level than STAT5b, as confirmed here by RNA-seq; furthermore, its loss has much less impact on hepatic expression of sex-biased genes (57). STAT1, although classically activated by interferon-γ stimulation (45), can also be activated by GH in liver (61). STAT1 is up-regulated in STAT5-KO liver (Fig. 2), perhaps as an adaptive response in the absence of competition by STAT5 for binding to their shared intracellular docking sites on GH receptor (28). STAT1 has a DNA-binding motif distinct from that of STAT5 (61) and consequently has unique target genes, and thus is unable to compensate for the loss of STAT5-regulated sex-biased gene expression (as was seen here) or for other phenotypic consequences of liver STAT5 deficiency, including impaired cell proliferation and fatty liver development (28). Although STAT5-KO dysregulated only ~2% of stringently sex-independent genes in liver (vs. 57% of sex-biased genes), at least some of the sex-independent genes and DMRs responsive to STAT5-KO may reflect adaptive mechanisms in the liver. Indeed, the sex-independent genes induced by STAT5-KO in both male and female liver are enriched for immune-specific genes (Fig. 2), at least some of which may be STAT1 targets.

In the *Alb*-Cre-induced hepatocyte-specific STAT5-KO model used here, STAT5 deletion occurs during late gestation, when the Cre transgene is activated by the *Alb* gene promoter (46). Therefore, the DNA methylation changes we observed may reflect late gestational or postnatal effects of STAT5 deficiency. Mammalian DNA methylation patterns are established at many sites during embryogenesis and then maintained throughout life (62, 63); however, DNA methylation continues to evolve postnatally at a subset of genomic regions in terminally differentiated cells (39, 40, 64). In mouse hepatocytes, demethylation at enhancer elements occurs after birth, when it is required to establish postnatal chromatin accessibility and gene transcription patterns (40). Our finding of enhancer hypermethylation in STAT5-KO liver indicates that STAT5 regulates demethylation of these genomic regions, which may contribute to chromatin opening and activation of enhancers critical for postnatal liver function. Our finding that these regions are also enriched for DNA motifs for other transcription factors important for sex-biased liver function (Fig. 8) suggests STAT5 may work with these other factors to recruit the TET demethylases (65) that actively demethylate these enhancer regions. Such demethylation events likely occur around puberty (39), when pituitary GH secretion becomes pulsatile in males, leading to intermittent activation of liver STAT5 (16, 17, 66), one of the key driving forces for the striking sex-differences in liver gene expression that become widespread at that time (67). Further study is needed to determine whether CpG demethylation of these enhancers by STAT5 is a permanent, irreversible event (39), or whether the methylation status of these enhancers is dynamically responsive to the repeated activation of STAT5 in male liver by plasma GH pulses, as was recently found for other sex-biased epigenetic events (16).

In contrast to STAT5-dependent enhancer DNA demethylation, we found that alteration of promoter DNA methylation is not a common mechanism for STAT5 regulation of sex-biased gene expression. Although CpG islands are frequently found at or nearby gene promoters, where they are mostly hypomethylated, even when the gene is transcriptionally silent (68), few promoters become activated in cells deficient in DNA methyltransferase activity, despite a 95% reduction in global DNA methylation and the creation of thousands of active enhancers (69–71). This suggests that DNA methylation does not drive promoter activation or repression, but rather, may be a more common regulatory mechanism in enhancers. The significance of DNA methylation at enhancer sites is apparent in cancers (69, 72, 73), where enhancer CpG methylation is more closely associated with changes in gene expression than is promoter methylation. Our finding that STAT5 primarily alters DNA methylation at enhancer elements is consistent with these findings, and is reminiscent of our earlier observation that sex-biased histone marks are more pronounced at distal sex-biased DHS than at sex-biased gene promoters (29).

STAT5 can respond to many extracellular cytokines and growth factor signals, including GH, and is the direct downstream effector of the hypothalamo-pituitary-liver GH axis. Liver STAT5 signaling is susceptible to disruption by xenobiotics, which can dramatically alter hepatic transcriptomic profiles and impact the ability of liver to metabolize foreign chemicals, increasing susceptibility to liver diseases, including metabolic disorders, fatty liver, hepatocellular carcinoma and obesity (4, 9). Our studies elucidating the interplay between STAT5-dependent DNA methylation and sex-biased gene expression have strong implications for understanding mechanisms underlying diseases associated with dysregulation of liver STAT5 function, which is a common occurrence (74, 75). Our findings suggest that STAT5 plays a critical role in shaping both sex-independent and sex-dependent enhancer elements via DNA methylation, which ultimately contributes to the control of proper RNA output for normal liver function. Given that DNA methylation is considered one of the more stable chromatin modifications, that once established, contribute to durable epigenetic memory through multiple cell divisions (76), dysregulation of STAT5 activity could lead to persistent effects on DNA methylation profiles at regulatory elements in the liver.

## Materials and Methods

### Liver tissue

Livers used for the analysis reported here were the same tissue samples used in our earlier work (42) and were originally obtained from Dr. Lothar Hennighausen (NIDDK, NIH). The livers were from 8 to 12-wk old hepatocyte STAT5-deficient males and females (KOM and KOF, respectively), and floxed controls (wild-type with respect to STAT5, i.e., WTM and WTF, respectively). The hepatocyte-specific STAT5-deficient mouse line (STAT5-KO) was developed by mating albumin promoter-driven Cre transgenic mice (FVB/N) with C57BL/6 x 129J mice having a floxed *Stat5a-Stat5b locus* (28). All mouse work was performed according to the Animal Research Advisory Committee Guidelines of NIH (https://oacu.oir.nih.gov/animal-research-advisory-committee-guidelines) and was approved by the NIDDK Animal Care and Use Committee.

### RNA-seq sample preparation and data analysis

A portion of each frozen liver was used for RNA extraction with TRIzol reagent (Invitrogen Life Technologies Inc., Carlsbad, CA) following the manufacturer’s instructions. Total liver RNA was isolated from n=5-6 individual livers per sex and genotype (WTM, WTF, KOM, KOF). Two RNA-seq libraries (biological replicates) were prepared for each group, with each sequencing library comprised of a pool of RNAs obtained from n=2-3 mouse livers. Libraries were prepared using the Illumina TruSeq RNA library preparation kit (Illumina, cat# RS-122-2001) starting with total liver RNA depleted of ribosomal RNA using NEBNext rRNA Depletion Kit. Paired-end sequence reads (50 nt reads) were generated on an Illumina HiSeq instrument at the New York Genome Center (New York, NY). RNA-seq data was analyzed using a custom pipeline developed in our lab (16). Briefly, FASTQ files with raw sequence reads were aligned to mouse genome build mm9 (NCBI 37) using Tophat (version 2.0.13) (77) using default parameters. FeatureCounts (78) was used to count sequence reads mapping to gene bodies within gene bodies of RefSeq genes, and EdgeR (79) was used to identify differentially expressed genes. 9,175 genes (**Table S1**) were identified as liver-expressed genes based on the criteria of >1 FPKM average expression in either male or female wild type mouse liver. An FDR < 0.05 was used to identify differentially expressed genes between male and female liver (sex-specific genes; 132 male-biased and 142 female-biased genes, based on WTM/WTF comparison) and for genes responsive to STAT5 loss (KOM/WTM, n=365 genes; and KOF/WTF, n=278 genes). Genes with FDR > 0.05 were deemed non-responsive to STAT5 loss (Table S1A). Stringent sex-independent genes (n=6,306) were identified by |fold-difference| < 1.2 and FDR > 0.1 for the WTM/WTF comparison.

### STAT5 ChIP-seq peaks nearby genes

Genomic coordinates of male-biased, female-biased and sex-independent STAT5 ChIP-seq peaks were downloaded from the Supplementary Materials of (12). STAT5 ChIP-seq peaks within 10 kb upstream or downstream to genes of interest were identified using the Bedtools (80) command: `bedtools intersect −c −w 10000` and are shown in Table S1A.

### Calculation of correlation

Gene expression fold change values and differentially methylated values contain both positive and negative values. When calculating correlations between two conditions, fold change and differentially methylated values were first converted to absolute values to avoid artificial overestimation of correlation. Linear regression was performed in R, and R^2^ and p-values are reported.

### Reduced representation bisulfite sequencing (RRBS)

RRBS libraries were prepared from frozen liver tissues as previously described, with modifications (41). Briefly, ~50 mg of frozen liver tissue was homogenized by vigorously pipetting up and down the sample in 370 μl TE buffer (10 mM Tris, 1mM EDTA, pH 8) containing 10 μl of proteinase K solution (10 mg/ml), and 20 μl of 10% (v/v) SDS was quickly added to achieve 0.5% SDS final concentration. The homogenate was incubated at 50°C for 2 hr. Genomic DNA (gDNA) was purified by extraction with 400 μl phenol chloroform with the addition of 32 μl of 5 M NaCl, using a large-bore pipet tip with gentle inversions and without vortexing. Intact gDNA was spooled from the aqueous phase following the addition of an equal volume pf 100% ethanol. The spooled gDNA was treated with RNase in PBS buffer (2 μl of 10 mg/ml RNase A in 298 μl PBS) for 1 hr at room temperature. The RNA-free gDNA was washed in ethanol and thoroughly resuspended in TE buffer for Qubit quantification of DNA, taking care to first sample a large volume (10 μl) and to then shear the DNA by vigorous pipeting to ensure that the subsequent 2 μl input for Qubit analysis is as homogenous as possible. 500 ng of intact gDNA was digested overnight with 2.5 μl of MspI (20,000 units/ml; NEB cat. R0106S) in the manufacturer’s supplied buffer (40 μl total volume) at 37°C, with the lid heated to 40°C to insure more complete digestion. The reaction was subjected to end-repair in the same reaction by adding 1 μl of Klenow fragment (New England Biolabs cat. M0210S, 5000 U/ml) and 1 μl of dNTP mix (10 mM dA, 1mM dG, 1 mM dC). Samples were then incubated at 30°C for 20 min, followed by 37°C for 20 min, and then purified by phenol chloroform extraction. 450 ng of MpsI-digested, end-repaired material (quantified by Nanodrop measurement) was ligated to Illumina adaptors supplied in the NEB DNA-seq kit (NEB cat. E7370S, using 1/20 diluted adaptor) and NEB Quick DNA ligase kit (NEB cat. E6056S) by incubation at 16°C overnight without a heated lid. After USER enzyme digestion and cleanup with Zymo’s DNA Clean and Concentrator-5 (Zymo cat. D4003T), the adaptor-ligated MspI fragments were bisulfite-treated for two consecutive rounds using Zymo’s EZ DNA Methylation-Gold Kit (Zymo cat. D5005) per the manufacturer’s instructions. A portion (5 μl) of each 12 μl bisulfite-treated DNA sample was amplified using the KAPA HiFi HS Uracil+ polymerase system (Kapa Biosytems/Roche cat. KM2800) for ~12-15 PCR cycles, with the optimal cycle number determined empirically to find the lowest number of cycles that yielded sufficient material without over-amplification. Finally, size selection of fragments 160-340 bp was accomplished by performing a double SPRI purification of 0.65x and 0.55x, as follows. SPRI beads (32.5 μl) were added to each 50 μl reaction (0.65x). After mixing well, the tubes were placed on magnetic rack until the beads were separated from the supernatant. The supernatant was transferred to fresh tubes, to which fresh SPRI beads (27.5 μl; 0.55x) were then added to the supernatant. The beads were washed twice in 200 μl of 80% ethanol solution, air dried for 5 min and eluted in 0.1X TE buffer. The final RRBS sequencing libraries were analyzed on a 1% agarose gel stained with SYBR green, quantified by Qubit analysis and sequenced on an Illumina HiSeq instrument (paired-end, 50 bp reads) with no more than ~30% of a sequencing lane being RRBS material. RRBS libraries were prepared and sequenced for the following samples with the indicated numbers of biological replicate livers: WT male (n=6), WT female (n=5), STAT5-KO male (n=6), and STAT5-KO female (n=6).

### RRBS data processing

About 15 million paired-end sequence reads were obtained for each RRBS sample. Raw FASTQ files were first processed by Trim Galore (version 0.4.3) using options: --rrbs --paired. The trimmed read1 and read2 files were mapped by Bismark (81) using default parameters. PileOMeth was used to extract and quantify all CpG dinucleotides in the bam files, outputting the location, total coverage, number of thymines and number of cytosines of each CpG in MethylKit-compatible format (parameter: --methylKit), while avoiding any double-counting of reads where paired-end reads for a given fragment overlap. The PileOMeth results were directly input into MethylKit, two sample groups at a time, for pair-wise differential analysis. MethylKit (82) differential analysis was performed at the level of both 100 bp tiles and individual CpGs using default parameters combined with the following: context=“CpG”, filterByCoverage (count=10), destrand=T. A total of 4 comparisons were performed in a Treatment vs. Control configuration, yielding differential methylation values (diff.meth), where the absolute value indicates the magnitude of change, and positive or negative signs indicate hypermethylation or hypomethylation of the treatment group relative to the control group, respectively. Tiles and individual CpGs with a significant change in DNA methylation were defined using a threshold fold-difference of 15% and q-value < 0.05, and were combined such that individual CpGs not captured at the tile level were included. Significant tiles and significant individual CpGs exhibit significant overlap with each other, and are summarized in **Table S2** (also see below). Here, differentially methylated regions (DMRs) refers to 100 bp genomic regions that fit one of the following criteria: 1) 100 bp tiles that were significantly differentially methylated at the level of overall tile statistics and also contain one or more CpGs that showed significant differences in methylation at the individual CpG level (this describes the majority of DMRs); 2) 100 bp tiles that were significant in their overall tile statistics but did not show statistically significant differences in individual CpGs; and 3) 100 bp regions that were not significantly differentially methylated at the level of overall tile statistics, but that contain at least one significant individual CpG, whose diff.meth values were averaged based on those CpGs that showed significant differences in methylation. The criteria by which the DMRs were identified is flagged in **Table S2**. In the majority of analyses, all DMRs were utilized.

A total 1,355,894 CpGs were captured by all 23 liver RRBS samples (Fig. S1, Supplemental file: all_cpG_covered_by_STAT5KO_23samples.txt). To generate this set of individual CpGs, raw PileOmeth files were de-stranded using a custom Python script, script27_destranding_pileometh.py; see Supplementary materials), which collapses CpG coordinates from opposite strands into a single coordinate, to facilitate downstream analyses, as follows. Each CpG is symmetrical in the genome, which means that each CpG will have two coordinates, e.g. chr1:100 (+) and chr1:101 (-). The de-stranding script aggregates the genomic position from the (+) strand and from the (-) strand into a single genomic position. Next, the frequency of occurrence of each CpG in each of the 23 RRBS-seq samples was determined, and only those CpGs that were covered by all 23 samples were considered. By segmenting the mouse genome into 100 bp non-overlapping windows, these 1,355,894 CpGs were aggregated into 232,882 windows (Supplemental file: complete_23STAT5_unique_tiles_unique.txt). The number of 100 bp windows (tiles) is lower than the number of individual CpGs because many windows contain multiple CpGs.

### Rationale for merging of tile and individual CpG statistics

As described above, DMRs were determined by MethylKit pair-wise comparison using a tile-based method and also an individual CpG method. The tile-based analysis gives higher statistical power due to the aggregation of signals from multiple CpGs within a 100 bp genomic region. For example, if a tile contains 3 CpGs, all of which are only slightly differential, the tile statistics may give a modest differential methylation value but with a low FDR, whereas individual CpG analysis would yield a high FDR for each CpG. In a different scenario, if a single CpG within a tile region is highly differentially methylated but is situated nearby other CpGs that are not differentially methylated, the signal from the highly differential CpG will be ‘diluted’ by the averaging used in the tile analysis, causing the loss of a significantly differential CpG. Because most tiles contain only one CpG, results of the tile and individual CpG analyses are similar for the most part. The final list of differential CpGs includes 2,375 significant tiles, as well as 1,514 100-bp genomic regions with individual CpGs that were captured by the individual CpG analysis but not by the tile analysis, for a combined total of 3,889 differential 100 bp genomic regions, is presented and further explained in Table S2.

An example of the R code tile and individual CpG differential analysis as follows:

~~~
#my.list refers to file locations for 11 PileOmeth output files
myobj_1 <-
methRead(my.list,sample.id=list(‘M1_WT’,’M2_WT’,’M3_WT’,’M4_WT’,’M5_WT’,’M6_WT’,’F1_WT’,’F2_WT’,’F3_WT’,’F4_WT’,’F5_WT’),assembly=‘mm9’,context=‘CpG’,treatment=c(1,1,1,1,1,1,0,0,0,0,0))
filtered.myobj_1<-filterByCoverage(myobj_1,lo.count=10,lo.perc=NULL,
hi.count=NULL, hi.perc=99.9)
meth_1<-unite(myobj_1,destrand=T)
tiles_1<-tileMethylCounts(meth_1,win.size=100,step.size=100)
#calculate differential 100 bp tiles
myDiff_tiles_1<-calculateDiffMeth(tiles_1)
#calculate differential individual CpG
myDiff_cpg<-calculateDiffMeth(meth_1)
~~~

### Generation of raw methylation heat maps

An intermediate file containing raw methylation values for all 100 bp tiles was generated by inputting Raw PileOMeth files into MethylKit and using the following command: `tileMethylCounts(pileometh_input,win.size=100,step.size=100)`. Raw methylation values were retrieved for only the differential tiles, which were visualized as a heat map using R. These raw methylation values in WTF, KOM and KOF samples were tested against WTM and evaluated using two-tailed Student t-test. For the heat maps shown in Fig. 3C, only DMRs determined by tile statistics with stringent threshold (diff.meth > 25%, qval < 0.05) were included. The heat map in Fig. 3C presents raw methylation values of DMRs that were determined to be significantly differential by tile analysis between STAT5-KO male liver and wild-type male liver when determined by tile analysis, i.e., excluding DMRs identified by individual CpGs only.

### DMRs in gene promoters

Gene promoters were defined as 5 kb upstream of the gene transcription start site. Thus, for genes on the + strand, the promoter coordinates were set as (start position – 5000, start position); for genes on the - strand, coordinates were set as (end position, end position + 5000). DMRs positioned in a promoter region (1 nt overlap) were identified using the Bedtools windows command (Table S3).

### Distribution of CpGs across chromatin states and annotated genomic features

To determine the distribution of chromatin states across All Tiles (see Fig. 5B), the 232,882 CpG-containing 100 bp tiles were mapped to 14 chromatin states previously defined in male mouse liver (29) using ChromHMM (83) and the Intersect command of Bedtools (80). Custom Bash commands and R scripts were used to calculate the frequencies of the 14 chromatin states in each bin and the results were plotted by ggplot. Chromatin state distributions were similarly determined for the sets of DMRs identified in the WTM/WTF, KOM/WTM and KOF/WTF comparisons, using the full set of DMRs (i.e., DMRs identified by tiles and DMRs identified by individual CpGs) described in Table S2 for each comparison. The relative frequency of occurrence of each chromatin state in each 100 bp bin across a 4 kb window surrounding each hypomethylated and hypermethylated DMR. To calculate enrichment scores, the 14 chromatin states were grouped into five super states, designated as inactive (states 1, 2, 3), enhancer (states 5, 6, 8, 9, 10), bivalent (state 12), transcribed (states 13, 14) and promoter (states 7, 8). Enrichment of DMRs for these five super states relative to ‘all CpGs’ (i.e., the set of 232,882 tiles covered by our RRBS samples) was calculated as follows: (number of differential DMRs in chromatin state-X / total number of differential DMRs) / (number of all DMRs in chromatin state-X / 232,882). Statistical significance was determined in R by two proportion Z-test using the *prop.test* function.

### TAD analysis

DMRs were grouped by their chromatin states defined in wild type mouse liver (29). DMRs and genes were then mapped to topologically associating domains (TADs) defined for mouse liver (51), and the overlap of DMR-containing TADs and gene-containing TADs was analyzed using datasets provided in Table S4. As a control, for each chromatin state, comparably sized sets of RRBS tiles that did not show significant sex differences or response to STAT5 loss were randomly sampled 50 times and mapped to TADs. The significance of DMRs occurring in TADs containing sex-biased, STAT5-responsive genes compared to the extent of overlap observed for the random controls was assessed by t-test. In addition, the distance from the start of the DMR to the TSS of sex-biased, STAT5 responsive genes was determined and analyzed using box plots, where DMRs that are > 10^6^ bp away from genes are deemed outliers and were excluded from analysis.

### Gene set enrichment analysis (GSEA)

DMRs determined by tile statistics with stringent threshold (qval < 0.05) were used. DMRs were ranked from high (hypermethylated) to low (hypomethylated). This list of ranked DMRs as well as the list of DMRs with nearby (within 10 kb) male-biased and female-biased DHS (52) (Table S5), were input into the GSEA desktop application for pre-ranked test analysis (84), with the number of permutations set to 1000.

### MEME DNA motif analysis

The Analysis of Motif Enrichment (AME) from the MEME suite (85) was used to assess the enrichment of the STAT5 motif in the tile sequences. The following command was used: ame −control random_sequences.txt −evalue-report-threshold 1 −o comp1_N comp1_N_sequence.txt STAT5_meme_motif.txt, where random_sequences.txt is a FASTA files of 100 bp sequences from randomly selected genomic regions that are not covered by our RRBS experiments; comp1_N_sequences.txt is the fasta file of the DMR tile sequences; and STAT5_meme_motif.txt is the MEME file of the STAT5 motif (see Supplemental materials).

### Cumulative frequency plots of STAT5 ChIP-seq peaks nearby DMRs

DMRs based on both tiles and individual CpG statistics for three comparisons (WTM/WTF; KOM/WTM; KOF/WTF) were separated into hypomethylated and hypermethylated DMRs. In addition, a file containing 1000 randomly selected non-differential methylated genomic regions was used. Sex-independent, male-biased and female-biased STAT5 ChIP-seq peaks were from (12). The analysis was based on 14,910 ChIP-seq peaks for STAT5 (see stat5b_merged_ss_sorted.txt, in Supplemental materials) from (12) after excluding 184 peaks on ChrX and ChrM from the full set of 15,094 ChIP-seq peaks reported in that study. Male-biased STAT5 binding occurs at 1,661 of the 14,910 sites, and female-biased binding occurs at 1,782 sites. The Bedtools command `bedtools closest −d −t` was used to find the distance to the nearest STAT5 peak for each DMR. For each of the seven sets of DMRs, DMRs were first ranked by distance to the nearest STAT5 peak from low to high, then cumulative values (which range from 0 to 1) were assigned to the ranked DMRs as follows: (position in ranked list / total number in list). The cutoff for the distance to nearest STAT5 peak was set at 10 kb. The distance (x axis) and cumulative frequency values (y axis) were plotted for all sets of DMRs.

### Homer DNA motif analysis

The Homer motif analysis tool (53) was used to find enriched motifs at DMRs relative to a background set comprised of a random selection of one thousand 100 bp regions that were detected by RRBS but not differentially methylated. Homer enrichment analysis of known motifs (Table S6) is based on 332 motifs in the HOMOR motif database that were mostly curated from published ChIP-seq datasets (http://homer.ucsd.edu/homer/motif/motifDatabase.html). The command for Homer motif analysis was as follows: *findMotifsGenome.pl input.txt mm9 output_folder −size 100 −bg random_1000.txt* where input.txt contains the genomic coordinates of DMRs to be analyzed for enriched regulatory elements; mm9 specifies the genome; “-size 100” indicates that the genomic regions in input.txt are 100 bp long; and “-bg random_1000.txt” specifies the coordinates for the background regions.

### Data availability

Raw and processed sequencing data are available at GEO (https://www.ncbi.nlm.nih.gov/geo/) under accession numbers GSE103885 (RNA-seq samples) and GSE103886 (RRBS samples).

## Acknowledgments

This work was supported in part by NIH grant DK121998 (to DJW). The funding agency had no role in study design, data collection and interpretation, or the decision to submit the work for publication. The authors thank Dr. Lothar Hennighausen, NIDDK, for originally providing the mouse liver tissue samples used in these analyses. The authors state they have no financial conflicts of interest to disclose.

## Author contributions

PH and DJW jointly conceptualized the project and designed the study. PH carried out all of the laboratory experiments, developed and implemented the methods for computational analysis, carried out data analysis and figure preparation. DJW obtained grant funding and supervised the study. PH and DJW jointly drafted the manuscript and DJW revised and edited the manuscript for publication.

